# Assessing the impact of artifact correction and artifact rejection on the performance of SVM-based decoding of EEG signals

**DOI:** 10.1101/2025.02.22.639684

**Authors:** Guanghui Zhang, Steven J. Luck

## Abstract

Numerous studies have demonstrated that eyeblinks and other large artifacts can decrease the signal-to-noise ratio of EEG data, resulting in decreased statistical power for conventional univariate analyses. However, it is not clear whether eliminating these artifacts during preprocessing enhances the performance of multivariate pattern analysis (MVPA; *decoding*), especially given that artifact rejection reduces the number of trials available for training the decoder. This study aimed to evaluate the impact of artifact-minimization approaches on the decoding performance of support vector machines.

Independent component analysis (ICA) was used to correct ocular artifacts, and artifact rejection was used to discard trials with large voltage deflections from other sources (e.g., muscle artifacts). We assessed decoding performance in relatively simple binary classification tasks using data from seven commonly-used event-related potential paradigms (N170, mismatch negativity, N2pc, P3b, N400, lateralized readiness potential, and error-related negativity), as well as more challenging multi-way decoding tasks, including stimulus location and stimulus orientation. The results indicated that the combination of artifact correction and rejection did not improve decoding performance in the vast majority of cases. However, artifact correction may still be essential to minimize artifact-related confounds that might artificially inflate decoding accuracy. Researchers who are decoding EEG data from paradigms, populations, and recording setups that are similar to those examined here may benefit from our recommendations to optimize decoding performance and avoid incorrect conclusions.

## 1. Introduction

The common view in electroencephalogram (EEG) research is that large artifacts caused by blinks and other sources (e.g., head movements, skin potentials, muscle contractions) can create both random noise and systematic differences across groups or conditions (reviewed by Luck, 2014). As a result, artifacts can lead to incorrect conclusions, either because of insufficient statistical power to detect real effects or because of confounds that produce bogus effects (Zhang, Garrett, Simmons, et al., 2024). To address this problem, some kind of artifact correction and/or artifact rejection approach is used in the vast majority of conventional event-related potential (ERP) studies. Guidelines for ERP studies therefore recommend including artifact correction and/or rejection as a part of the preprocessing pipeline (Duncan et al., 2009; Keil et al., 2014, 2022).

Recently, *pattern classification* or *decoding* approaches based on machine learning (ML) have received growing interest in EEG and ERP research. A major reason for this is that decoding analyses can distinguish subtle differences between experimental stimulus classes that cannot be detected by traditional univariate approaches (Hebart & Baker, 2018; Peelen & Downing, 2023). For example, a greater effect size can be obtained for decoding analyses than for traditional univariate analyses across a broad range of commonly-used ERP paradigms (Carrasco et al., 2024). In addition, multivariate decoders may be able to disregard noisy channels while retaining meaningful information (Ashton et al., 2022; Carlson et al., 2019; Grootswagers et al., 2017).

Of course, noise may still degrade decoding performance. Noise from ocular artifacts may be particularly problematic, because blinks and eye movements are large signals that are typically present on only a subset of trials, creating substantial trial-to-trial variance. This trial-to-trial variance may be particularly deleterious for decoding, which depends on a cross-validation procedure in which the decoder is trained on one subset of trials and then tested on a new subset of trials; trial-to-trial variation in the signals may cause the training and test sets to differ, reducing decoding accuracy. On the other hand, ocular artifacts may also artificially inflate decoding accuracy. That is, blinks or eye movements may differ systematically across the classes being decoded, leading to above-chance classification even if the neural signals are identical across classes (Hong et al., 2020). To minimize both nonsystematic artifacts (noise) that might degrade decoding and systematic artifacts that might inflate decoding, artifact correction and/or artifact rejection preprocessing steps are commonly employed prior to decoding.

There are two major approaches to dealing with artifacts, namely artifact rejection and artifact correction. In artifact rejection, each trial is *marked* as containing or not containing a problematic artifact, and the marked trials are excluded from subsequent analyses. This may reduce noise by eliminating trials with large non-neural signals, and it may also reduce the confounding effects of systematic artifacts, but it has the downside of decreasing the number of trials available for decoding. However, the reduction in noise resulting from artifact rejection can outweigh the reduction in the number of trials available for analysis in conventional ERP analyses (Zhang, Garrett, Simmons, et al., 2024), and many previous decoding studies have applied artifact rejection prior to classification (e.g., Alizadeh et al., 2017; Despouy et al., 2020; Mares et al., 2020).

An alternative approach is artifact correction, in which a method such as independent component analysis (ICA) is used to separately estimate the artifactual and non-artifactual sources that are combined in the measured signals and then reconstruct the EEG from the non-artifactual components. This does not reduce the number of trials available for analysis, so it has been used instead of artifact rejection in many decoding studies (e.g., Bae, 2021a, 2021b; Bae & Luck, 2018; Meier et al., 2022; Tu et al., 2023).

However, ICA-based artifact correction is appropriate only for artifacts that have a consistent scalp distribution (e.g., blinks and eye movements), and it is inappropriate for artifacts that vary in scalp distribution across trials (e.g., idiosyncratic movement artifacts). Thus, some decoding studies have used a combination of both artifact correction and artifact rejection (e.g., Den Ouden et al., 2023; He et al., 2024; Jaatinen et al., 2023; Kato et al., 2022; Q. Li et al., 2022; Y. Li et al., 2022; Xiong et al., 2024). This combination of correction and rejection has been found to be effective at both maximizing data quality and minimizing confounds in conventional ERP analyses (Zhang, Garrett, Simmons, et al., 2024). However, to our knowledge, there has not been a broad and systematic examination of whether these approaches to minimizing artifacts can enhance the performance of decoding analyses in a way that generalizes across a wide range of EEG/ERP datasets and classification tasks.

### 1.1 The goal of the present study

Several studies have explored the effectiveness of artifact minimization approaches on decoding performance from individual datasets with a relatively small number of participants (Bae, 2021a; Bae & Luck, 2019; Hong et al., 2020). However, it is not known whether the results of these studies will generalize to studies with different experimental paradigms, electrode densities, and numbers of stimulus classes.

The goal of the present study was to evaluate the impact of artifact minimization approaches on decoding performance in a systematic and generalizable manner. This was achieved by using a broad range of EEG/ERP experimental paradigms with varying experimental classes and different numbers/densities of electrodes. Previous research has indicated that support vector machines (SVMs) outperformed other classification algorithms, such as linear discriminant analysis and random forest, in simple ERP decoding analyses (Trammel et al., 2023). We therefore employed the SVM algorithm in the current study.

Our analyses included data from the publicly available ERP CORE (Compendium of Open Resources and Experiments; Kappenman et al., 2021) dataset, which includes six common ERP paradigms that can be used to isolate seven distinct ERP components: N170, mismatch negativity (MMN), P3b, N400, error-related negativity (ERN), N2pc, and lateralized readiness potential (LRP). In each of these paradigms, we performed binary decoding of the primary experimental classes (e.g., target versus nontarget for the P3b component, standard versus deviant for MMN component). Note that these datasets were collected from 32 channels in highly cooperative college students.

We also examined three additional datasets in which the differences between stimulus classes are not easily detectable using conventional univariate methods. The first two paradigms tasked participants with perceiving and remembering one of 16 distinct orientations presented on each trial (Bae & Luck, 2018). We conducted decoding analyses to determine which of the 16 orientations was presented, both during perceptual processing (100-500 ms) and working memory maintenance (500-1000 ms). Additionally, in one of these datasets, stimuli were presented at 16 different locations, allowing us to decode stimulus location as well as stimulus orientation. As in the ERP CORE dataset, the data from these two experiments were recorded from highly cooperative college students. To determine whether our results generalize to the broader population, we performed orientation decoding in an additional dataset with a similar orientation memory task but with data collected from a broad community sample of adults. This dataset included 64 channels.

Thus, our analyses included both 32-channel and 64-channel datasets, both simple binary classification tasks and complex multi-way classification tasks, and both college student and community samples. This makes it possible to draw relatively general conclusions about the impact of artifact rejection and correction on EEG pattern classification performance. Of course, these conclusions may not extend to vastly different experimental paradigms, participant populations (e.g., infants), recording hardware (e.g., saline-based electrodes), and recording environments (e.g., hospitals).

The present study focused solely on the extent to which artifact rejection and correction lead to improved decoding accuracy by decreasing artifact-related noise, not on whether they successfully eliminate confounds due to systematic artifacts (e.g., difference in blinking between different stimulus classes). We have previously developed an easy-to-implement method to assess whether such confounds have been successfully eliminated (Zhang, Garrett, Simmons, et al., 2024). Hong et al. (2020) also developed a simple approach to determining whether ICA has eliminated blink-related confounds. We focus on removing blinks with ICA, because they are the most ubiquitous large artifacts that can easily be corrected using ICA. We use artifact rejection rather than artifact correction to deal with artifacts that cannot easily be handled by ICA (e.g. idiosyncratic movement artifacts).

## 2. Methods

The analyses were performed on publicly available EEG datasets. The following sections briefly described the stimuli, tasks, recording methods, and preprocessing methods for each dataset; detailed descriptions can be found in the original papers. We will refer to the datasets as *ERP CORE* (Kappenman et al., 2021; data available at https://doi.org/10.18115/D5JW4R), *Orientations–1 and Orientations-2* (Bae & Luck, 2018), and Orientations-Community (unpublished study; data available at https://osf.io/s24de/). All the studies were approved by the Institutional Review Board at each data collection site, and all participants provided informed consent.

We carried out the data analyses within the MATLAB 2021b (MathWorks Inc) environment using EEGLAB Toolbox v2024.0 (Delorme & Makeig, 2004) and ERPLAB Toolbox v10.10 (Lopez-Calderon & Luck, 2014). All data and scripts are available at https://osf.io/s24de/.

### 2.1 Participants

The participants for the ERP CORE and Orientations-1/2 experiments were college students at UC Davis (*N* = 40 for ERP CORE; *N* = 15 for Orientations-1, *N* = 16 for Orientations-2). The participants for the Orientations-Community experiment (N = 27) were recruited from the Baltimore, Maryland community to serve as control subjects for a study of a clinical population (not analyzed here). We expected that the data would be noisier in this community sample than in the college students.

### 2.2 Experimental paradigms

Each of the six tasks in the ERP CORE lasted approximately 10 minutes, and each participant completed all six tasks within a single session. The N170 component was elicited using a face perception paradigm (Figure 1a). On each trial, participants were presented with one of four stimulus categories (face, car, scrambled face, or scrambled car) at the center of the display. Participants were instructed to distinguish between “texture” stimuli (scrambled face or scrambled car) and “objects” (face or car) by pressing one of two buttons. Only data from the face and car stimuli were analyzed in the present study. The mismatch negativity (MMN; Figure 1b) was elicited using a passive auditory oddball task.

**Figure 1.**
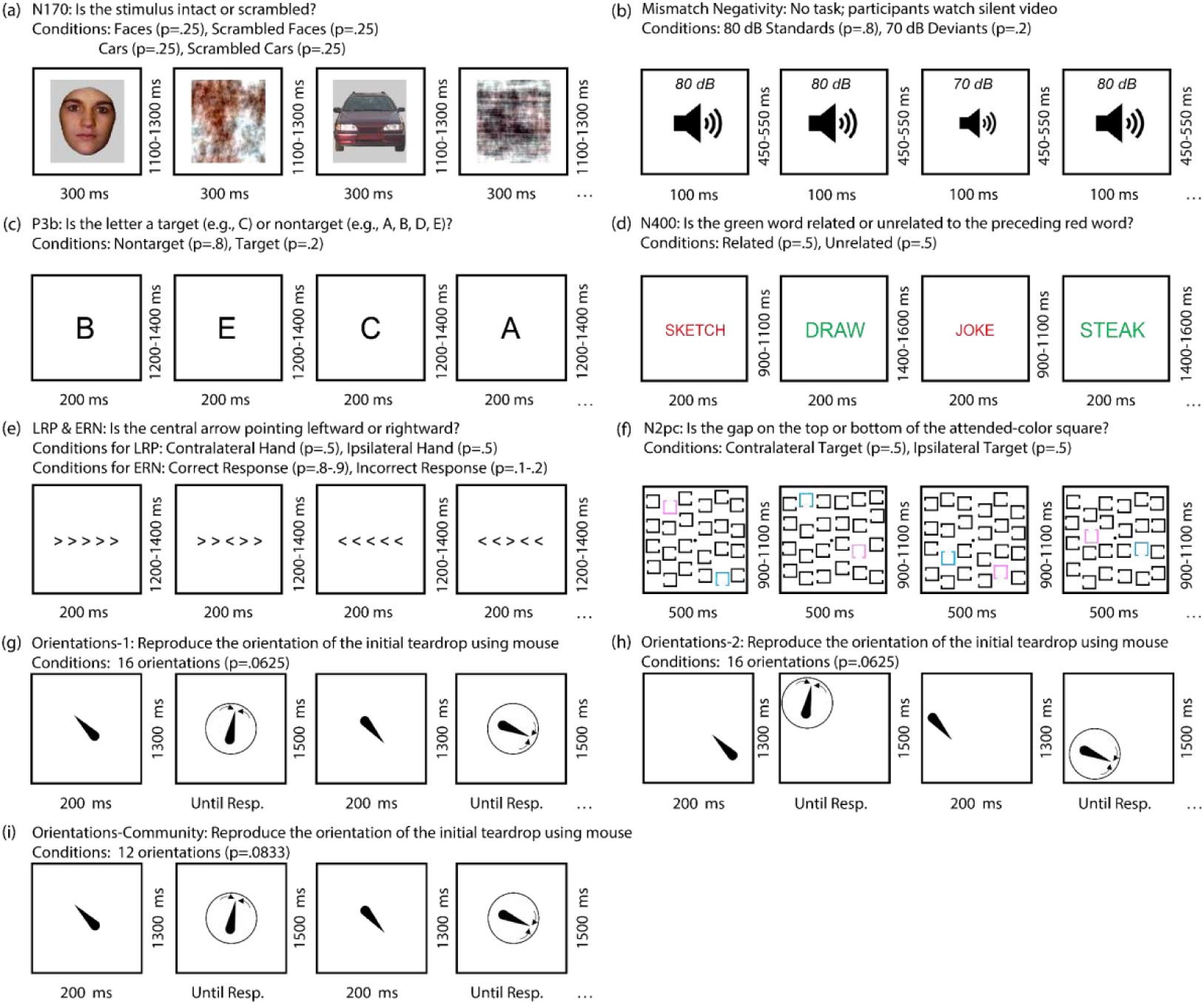
Example trials from the experimental paradigms. (a) N170 paradigm: Participants identified whether a given stimulus was intact (faces or cars) or scrambled (scrambled faces or cars). The present study focused exclusively on the unscrambled face and car trials. (b) Mismatch negativity paradigm: Participants watched a silent video while task-irrelevant standard tones (80 dB, probability =.8) and deviant tones (70 dB, probability =.2) were presented. (c) P3b paradigm: The letters A, B, C, D, and E were presented in random order, each with a probability of.2. One letter was designated as the target for each trial block, and participants identified whether a given stimulus was the target or a nontarget. (d) N400 paradigm: Participants determined whether a green target word was semantically related or unrelated to a preceding red prime word. (e) Flankers task for eliciting the lateralized readiness potential (LRP) and error-related negativity (ERN). Participants reported the direction of the central arrow (leftward or rightward) while ignoring the surrounding arrows. (f) N2pc paradigm: For each trial block, either pink or blue was designated as the target color. Participants reported whether the gap on the target-colored square was at the top or bottom of the square. (g) Orientations-1 paradigm: Participants memorized the orientation of an initial teardrop, retained it over a 1300-ms delay period, and then used a mouse to adjust the orientation of a test teardrop to match the original. (h) Orientations-2 paradigm: Identical to Orientations-1, except the teardrop appeared randomly at one of 16 different locations. (i) Orientations-Community: Identical to Orientations-1, except only 12 orientations were presented.

Participants listened to a series of standard stimuli at 80 dB (p=.8) and deviant stimuli at 70 dB (p =.2) while watching a silent video, with instructions to disregard the auditory stimuli. The P3b component (Figure 1c) was elicited using an active visual oddball task. Participants viewed a randomized sequence of letters (A, B, C, D, and E; p =.2 for each). One of these letters served as the target for a block of trials (e.g., A was the target and the others were non-targets for one block). Participants were directed to press one button for the target stimulus (p =.2) and a different button for any of the non-target stimuli (p =.8). The N400 component (Figure 1d) was elicited using a word pair judgment task. Each trial included a red prime word followed by a green target word; participants indicated whether the target was semantically related to the prime by pressing one of two buttons. The lateralized readiness potential (LRP) and the error-related negativity (ERN) were elicited using an Eriksen flankers task (Figure 1e). Participants identified the direction of a central arrowhead surrounded by congruent or incongruent arrowheads by pressing corresponding buttons with their left or right hand. The N2pc component (Figure 1f) was elicited using a visual search task. Participants identified the location (top or bottom) of a gap in squares of a designated color (pink or blue) by pressing buttons corresponding to each option. Throughout all tasks, participants-maintained fixation on a central point as instructed.

The three Orientations datasets came from slightly different variations of the same paradigm, administered to separate groups of participants. In the Orientations-1 paradigm (Figure 1g), a teardrop shape (the sample stimulus) appeared for 200 ms at the center of the display at the beginning of the trial. There were 16 potential teardrop orientations (at 22.5° intervals). Following a 1300-ms blank delay, participants were tasked with reproducing the orientation of the initial teardrop by using a mouse to adjust the orientation of a test teardrop. In the Orientations-2 paradigm (Figure 1h), the sample teardrop was again presented at one of 16 randomly chosen orientations, but the location of the teardrop varied across 16 locations around a circle of possible locations. The orientation and location of the teardrop varied independently. Participants were again tasked with reproducing the orientation after the delay period, but the location of the teardrop was task-irrelevant, and the test teardrop appeared at a randomly selected position. The Orientations-Community paradigm was identical to the Orientations-1 paradigm, except that the number of orientations was reduced from 16 to 12 (at 30° intervals).

### 2.3 EEG recording and preprocessing

In the ERP CORE paradigms, the EEG was recorded using a Biosemi ActiveTwo recording system (Biosemi B.V.) at a sampling rate of 1024 Hz. A fifth-order sinc antialiasing filter with a half-power cutoff at 204.8 Hz was applied during recording. The EEG was recorded from 30 scalp sites, including FP1/2, Fz/3/4/7/8, FCz/3/4/, Cz/3/4/5/6, CPz, Pz/3/4/7/8, PO3/4//7/8/9/10, Oz/1/2. Additionally, electrooculogram (EOG) electrodes were positioned adjacent to the eyes and below the right eye to monitor eye movements.

In the three Orientations paradigms, the EEG was recorded using a Brain Products actiCHamp recording system (Brain Products GmbH) at a 500 Hz sampling rate. A cascaded integrator-comb antialiasing filter with a half-power cutoff at 130 Hz was employed during recording. The recordings included the same EOG electrode locations used in the ERP CORE dataset along with the left and right mastoids. The Orientations-1 and Orientations-2 datasets included recordings from 27 channels (FP1/2, Fz/3/4/7/8, Cz/3/4, Pz/3/4/5/6/7/8/9/10, POz/3/4/7/8, Oz/1/2), and 57 scalp sites were included for the Orientations-Community dataset (FP1/2, AFz/3/4/7/8, Fz/1/2/3/4/5/6/7/8, FCz/1/2/3/4/5/6, Cz/1/2/3/4/5/6, CPz/1/2/3/4/5/6, TP7/8, Pz/1/2/3/4/5/6/7/9/10, POz/3/4/7/8, Oz/1/2).

Here we provide a brief overview of the key processing steps for the ERP CORE data; comprehensive descriptions can be found in the original paper (Kappenman et al., 2021).

The continuous EEG signal was resampled to 256 Hz. The EEG was referenced to the mean of the P9 and P10 electrodes (close to the left and right mastoids), except in the N170 paradigm where the average of all scalp sites served as the reference. A noncausal Butterworth filter was then applied (bandpass = 0.1 – 30 Hz, 12 dB/oct roll-off). ICA was then applied to correct the data for blinks and eye movements (see details in Section 2.4). The ICA-corrected data were subsequently epoched and baseline-corrected using the time windows delineated in Table 1. Bad channels were addressed through interpolation using EEGLAB’s spherical spline interpolation algorithm. Trials were marked for rejection if they contained blinks that would have prevented perception of the stimulus, extreme values in any channel, or erroneous behavioral responses (see details in Section 2.4).

**Table 1.** Epoch window, baseline period, electrode site, and measurement window used for each experiment.

| Experiment | Epoch (ms) | Baseline Period (ms) | Measurement window (ms) | # of Subjects | # of classes (M) | # of channels |
| --- | --- | --- | --- | --- | --- | --- |
| N170 | -200 to 800 | -200 to 0 | 110 to 150 | 36 | 2 (faces vs. cars) | 28 |
| MMN | -200 to 800 | -200 to 0 | 125 to 225 | 38 | 2 (deviants vs. standards) | 28 |
| P3b | -200 to 800 | -200 to 0 | 300 to 600 | 33 | 2 (targets vs. nontargets) | 28 |
| N400 | -200 to 800 | -200 to 0 | 300 to 500 | 36 | 2 (related vs. unrelated) | 28 |
| ERN | -600 to 400 | -400 to -200 | 0 to 100 | 35 | 2 (correct vs. error) | 28 |
| LRP | -800 to 200 | -800 to -600 | -100 to 0 | 37 | 2 (left hand vs. right hand) | 28 |
| N2pc | -200 to 800 | -200 to 0 | 200 to 275 | 35 | 2 (left target vs. right target) | 28 |
| Orientations-1 | -500 to 1500 | -500 to 0 | 100 to 500 or 500 to 1000 | 15 | 16 (16 orientations) | 27 |
| Orientations- 2 | -500 to 1500 | -500 to 0 | 100 to 500 or 500 to 1000 | 15 | 16 (16 orientations or 16 locations) | 27 |
| Orientations-Community | -500 to 1500 | -500 to 0 | 100 to 500 or 500 to 1000 | 27 | 12 orientations | 59 |

The same preprocessing steps were used for the Orientations datasets, except that the data were downsampled to 250 Hz and referenced to the average of the mastoids, and no trials were rejected because of behavioral errors (see Bae & Luck, 2018).

### 2.4 General artifact correction and rejection methods

ICA served as the method for artifact correction, implemented through the EEGLAB *runica()* routine with the infomax algorithm. This algorithm is widely utilized for artifact correction, being the default setting in EEGLAB (Delorme & Makeig, 2004). We used the default parameters because our aim was to assess the effectiveness of a commonly employed artifact minimization approach rather than trying to determine the optimal approach (which is beyond the scope of this study).

Current recommendations for ICA (Dimigen, 2020; Klug & Gramann, 2021; Luck, 2022) were not followed in the original preprocessing of these datasets, so we re-ran the ICA decomposition procedure for each dataset using the current recommendations. The new decomposition involved making a copy of the original continuous dataset for each participant, optimizing this parallel dataset for ICA, performing the ICA decomposition on this parallel dataset, and then transferring the ICA weights back to the original dataset. In this parallel dataset, a noncausal Butterworth band-pass filter was applied with half-amplitude cutoffs at 1 Hz and 30 Hz (a roll-off of 12 dB/octave) to minimize idiosyncratic noise that might otherwise interfere with the ICA decomposition^1^. We then resampled the data at 100 Hz to expedite the decomposition process. Finally, we removed break periods and segments with non-biologically plausible outlier voltages from the continuous EEG data. These segments, containing idiosyncratic noise, were identified and eliminated using the ERPLAB *pop_erplabDeleteTimeSegments()* function for break periods that were defined as intervals of at least 2 seconds without an event code, and the ERPLAB *pop_continuousartdet()* function for segments where the peak-to-peak amplitude within a specified window length exceeded a predetermined threshold. The window length and threshold were individually determined through visual inspection in the original ERP CORE resource, with window lengths ranging from 500 to 2000 ms and thresholds from 350 to 750 µV. The ICA decomposition was then performed on the resulting data.

In the ICA decomposition, all EEG and EOG electrodes were included, excluding bipolar channels and any “bad” channel necessitating interpolation, as identified in the original ERP CORE dataset. To identify independent components (ICs) related to blink artifacts, we utilized ICLabel, an automated IC classification system trained on numerous datasets with manually labeled ICs (Pion-Tonachini et al., 2019). ICLabel assigns a probability to each IC reflecting the likelihood that it reflects a specific artifact. We categorized an IC as representing blinks if ICLabel attributed it a probability of at least 0.9 for blink activity. For ICs with lower probabilities, indicating ambiguity, we visually assessed the time-course alignment with the VEOG-bipolar signal. When a strong alignment was observed, we classified the IC as a blink. While this semi-automatic classification method is common, it relies on expert subjective judgment and may not be fully reproducible. In a previous study (Zhang, Garrett, Simmons, et al., 2024), we also conducted analyses using fully automatic classification (i.e., without visual inspection), employing probability thresholds of 0.9 and 0.8. All three approaches yielded statistically identical results in almost every case (see the details in Zhang, Garrett, Simmons, et al., 2024). The results presented here are based on the semi-automatic approach.

Following the ICA decomposition on the heavily filtered parallel dataset, we transferred the IC weights to the original dataset, which had undergone minimal filtering. We then reconstructed the EEG using the non-blink ICs. This approach enabled us to utilize ICA weights derived from heavily filtered data, unaffected by idiosyncratic noise, while ensuring that these weights were applied to the original dataset, thus preserving the temporal resolution of the data.

Artifact detection and rejection were performed after artifact correction and after the data were epoched and baseline-corrected using a time range between the start of the baseline period and the end of the measurement time window (listed in Table 1). For example,-200 to 0 ms was used for P3b baseline correction and 300 to 600 ms was used for P3b component measurement, so we used a time window of-200 to 600 ms to detect and reject artifact-related trials. We explored several alternative approaches, such as rejecting trials with artifacts across the entire epoch (e.g.,-200-800 ms for P3b) or specifically within the time window of interest for one ERP component (e.g., 300-600 ms for P3b). These additional approaches produced similar results to the approach used here.

In addition to artifact correction, we systematically explored multiple algorithms and parameters to reject artifacts that remained after the ICA-based blink correction. In all cases, a given algorithm was used to mark epochs that exceeded a threshold (artifact *detection*), and the marked trials were excluded from the decoding analyses (artifact *rejection*).

We first examined a widely-used approach in which simple algorithms were used to mark trials that contained extreme values in any channel. Specifically, we applied ERPLAB’s simple voltage threshold (SVT) algorithm and moving window peak-to-peak (MWP) algorithm to the baseline-corrected data during the rejection window. The SVT algorithm marks epochs in which the absolute value of the voltage exceeds a threshold. The MWP algorithm slides a 200-ms moving window across the artifact window, computing the peak-to-peak amplitude within each moving window; the result is the maximum of the amplitudes across all the moving windows. For both algorithms, we explored multiple thresholds (see Section 2.5). For any given analysis run, the threshold was the same for both SVT and MWP (e.g., 100 µV for both algorithms). Note that a given epoch was excluded if an artifact was detected by either algorithm in any channel.

We separately tested an alternative artifact detection approach in which epochs were marked for rejection if they contained values that were statistically improbable in any channel. In this approach, we first took the single-sample voltage values across all participants, all channels, and all time points within the rejection window and computed a probability density function that quantified the probability of any given voltage value.

Voltages that are far from the mean are considered statistically improbable. We therefore flagged epochs in which the voltage at any channel within the rejection window exceeded a threshold that was based on the standard deviation (SD) of the probability density function. We tested thresholds ranging from 2 to 6 SDs.

### 2.5 Combining artifact correction and rejection

We tested four different combinations of artifact correction and artifact rejection. In an approach that we labeled *Baseline*, we did not apply either artifact correction or artifact rejection. We simply segmented and baseline-corrected the filtered EEG using the time windows shown in Table 1. All epochs were then used for decoding (except epochs with incorrect behavioral responses in ERP CORE experiments).

For the second approach, labeled *ICA-Only*, ICA was used to correct EEG data and no rejection of epochs was performed (except those with incorrect behavioral responses in ERP CORE experiments).

For the third approach, labeled *ICA+AbsThreshold*, we first applied ICA to correct blinks, and then we rejected epochs with voltage deflections that exceeded an absolute threshold in any channel as determined by the SVT and MWP algorithms from ERPLAB Toolbox. We systematically tested several different thresholds (50, 60, 70, 80, 90, 100, 200, 400 µV for ERP CORE experiments; 70, 80, 90, 100, 200, 400 µV for Orientations-1/2; 150, 200, 250, 300, 400, 500 µV for Orientations-Community). Note that a lower threshold will remove more artifacts but will also leave fewer trials available for decoding; by systematically varying the threshold, we could examine how the tradeoff between these two factors impacted decoding accuracy. We used different sets of thresholds for the different datasets because some of the thresholds that were appropriate for some datasets would have yielded too few trials for decoding for many of the participants in other datasets. Note that the goal was not to determine a single optimal threshold for rejection, which likely differs considerably across subject populations, but instead to ask in general how the benefits of rejecting trials in terms of noise reduction balances with the cost in terms of the number of trials available for decoding.

The fourth approach, labeled *ICA+StatThreshold*, was identical to the *ICA+AbsThreshold* approach except that we used a *statistical* rejection threshold based on the SD from the probability density algorithm. That is, we detected and rejected trials that contained voltages beyond a certain number of SDs away from the mean in the distribution of voltages across all subjects. We systematically varied the SD threshold for rejection (2, 3, 4, 5, and 6 SDs) just as we varied the threshold for rejection in the *ICA+AbsThreshold* approach.

### 2.6 SVM-based decoding methods

For the ERP CORE paradigms, we decoded which of the two stimulus classes was present for a given paradigm (see Table 1). For example, in the P3b experiment, we classified whether each stimulus was the target or a nontarget. For the Orientations-1 and Orientations-2 paradigms, we decoded which of the 16 orientations was present (collapsing across stimulus locations in Orientations-2). Additionally, in the Orientations-2 paradigm, we decoded which of the 16 locations contained the stimulus (collapsing across stimulus orientations). For the Orientations-Community paradigm, we decoded which of the 12 orientations was present. Thus, the number of classes to be decoded was 2 for each of the ERP CORE paradigms, 16 for the Orientations-1 and Orientations-2 paradigms, and 12 for the Orientations-Community paradigm.

We decoded the data separately at each time point for each participant. The MATLAB function *fitcsvm()* was applied to train the SVM when decoding 2 classes. When we decoded more than 2 classes, we employed the MATLAB function *fitcecoc()* to implement the error-correcting output codes approach (see Bae & Luck, 2018 for a detailed overview of this approach). For both procedures, the MATLAB function *predict()* was used to test the decoder.

We used a leave-one-out 3-fold cross-validation approach to avoid overfitting, as applied in previous studies (Carrasco et al., 2024; Zhang et al., 2024). For each decoding analysis, we divided the matrix for a given participant into separate matrices for each of the *M* stimulus classes (e.g., where *M* is 16 for the Orientations-1/2 experiment). To maximize decoding accuracy, we performed decoding on averaged ERPs, with one averaged ERP waveform for each of the 3 folds. That is, 3 averaged ERPs were created for each class, based on different random subsets of the trial. For example, in the Orientations-1/2 experiments, there were 16 stimulus classes with 40 trials for each stimulus when no artifact rejection was applied. We created 3 averages for each of the 16 orientations, with 13 randomly selected trials in each average. When artifact rejection was performed, fewer trials were available, and we reduced the number of trials per average as needed, but we ensured that the number of trials per average was equated across classes (because an unequal number of trials can spuriously inflate decoding accuracy).

In each participant, two of the three averages in each class were used to train the decoder and the one left-out average for each class was used to test the decoder. This gave us 16 test cases for the Orientations 1-/2 experiments, 12 test cases for the Orientations-Community experiment, and 2 test cases for each of the ERP CORE experiments. This process was repeated three times, each time using two different training cases and the remaining test case for each class. We then repeated this process 100 times, using a different random assignment of trials to averages for each iteration. Decoding accuracy was defined as the proportion of test cases that were correctly classified. The chance level was set at 1 divided by the number of stimulus classes (1/2 or 0.5 for the ERP CORE experiments, 1/16 or.0625 for the Orientations-1/2 experiments, and 1/12 or.083 for the Orientations-Community experiment).

### 2.6 Time-window averaging, effect size calculation, and statistical testing

To easily compare the decoding results for different artifact minimization approaches, we averaged the decoding accuracy across a predefined time window for each paradigm as shown in Table 1. For the ERP CORE experiments, we simply used the time window that was defined by Kappenman et al. (2021) for the conventional univariate analysis for a given ERP component. For the other paradigms, which involved working memory tasks, we computed the decoding accuracy in both a perceptual time window (100–500 ms) and a working memory time window (500–1000 ms). Paired *t* tests were used to compare decoding accuracy for each artifact approach with the baseline approach in which no artifact correction or rejection was performed.

For engineering applications, the best decoding method is usually defined as the one that provides the highest decoding accuracy. In scientific applications, however, the main goal is typically to determine whether the mean decoding accuracy across participants differs from chance, which depends on both the mean decoding accuracy and the variability across participants. Therefore, it is beneficial to consider Cohen’s *d_z_* metric of effect size (Lakens, 2013), which accounts for both the mean decoding accuracy and the variability among participants. To this end, we computed Cohen’s *d_z_* for each analysis by subtracting the chance level from the mean decoding accuracy for a defined time-window across participants and dividing by the standard deviation across participants. We used bootstrapping with 10,000 iterations to estimate the standard error of the *d_z_* value.

### 2.7 Quantifying Noise

Given that the cross-validation procedure involves training the decoder on one subset of trials and testing it with a different set of trials, any noise that increases the trial-to-trial variability will directly decrease decoding accuracy. This was one of the reasons we included the Orientations-Community dataset, which we assumed would be noisier than the other datasets. To quantify the noise level for each dataset, we computed the standard error of the mean for each participant’s averaged ERPs (including all trials) for each combination of, channel, condition, and time point (see Luck et al., 2021, for an explanation of why the standard error is a particularly good metric of noise for averaged ERPs). To compute a single value for each participant, we calculated the root mean square of the standard errors across time points, electrode sites, and conditions. The results are summarized in Supplementary Table S1.

## 3. Results

### 3.1 Simple binary decoding cases

In this section, we describe the binary decoding results for the ERP CORE experiments. Note that Kappenman et al. (2021) found that there was a statistically significant difference between the two experimental conditions for each ERP component during the measurement window when the data were analyzed using a conventional univariate analysis, so we do not include any univariate analyses here.

Figure 2 shows the results of our decoding analysis, in which we attempted to decode which of the two stimulus or response categories defined each of the seven ERP components (e.g., faces versus cars for the N170 component, correct versus incorrect for the ERN). The left column shows mean decoding accuracy across the different artifact minimization approaches, and the middle column shows the corresponding effect sizes (Cohen’s *d_z_*), which take into account both mean and the across-subject variability of decoding accuracy. The ICA-Only and Baseline approaches yielded nearly identical decoding accuracy and effect sizes for the N170, P3b, N400, N2pc, and LRP analyses. For the MMN, both decoding accuracy and *d_z_* were increased somewhat by the ICA-Only approach compared to the Baseline approach, but this increase was not statistically significant. For the ERN, decoding accuracy and *d_z_* were reduced somewhat for the ICA-Only approach relative to the Baseline approach, but this reduction was not statistically significant.

**Figure 2.**
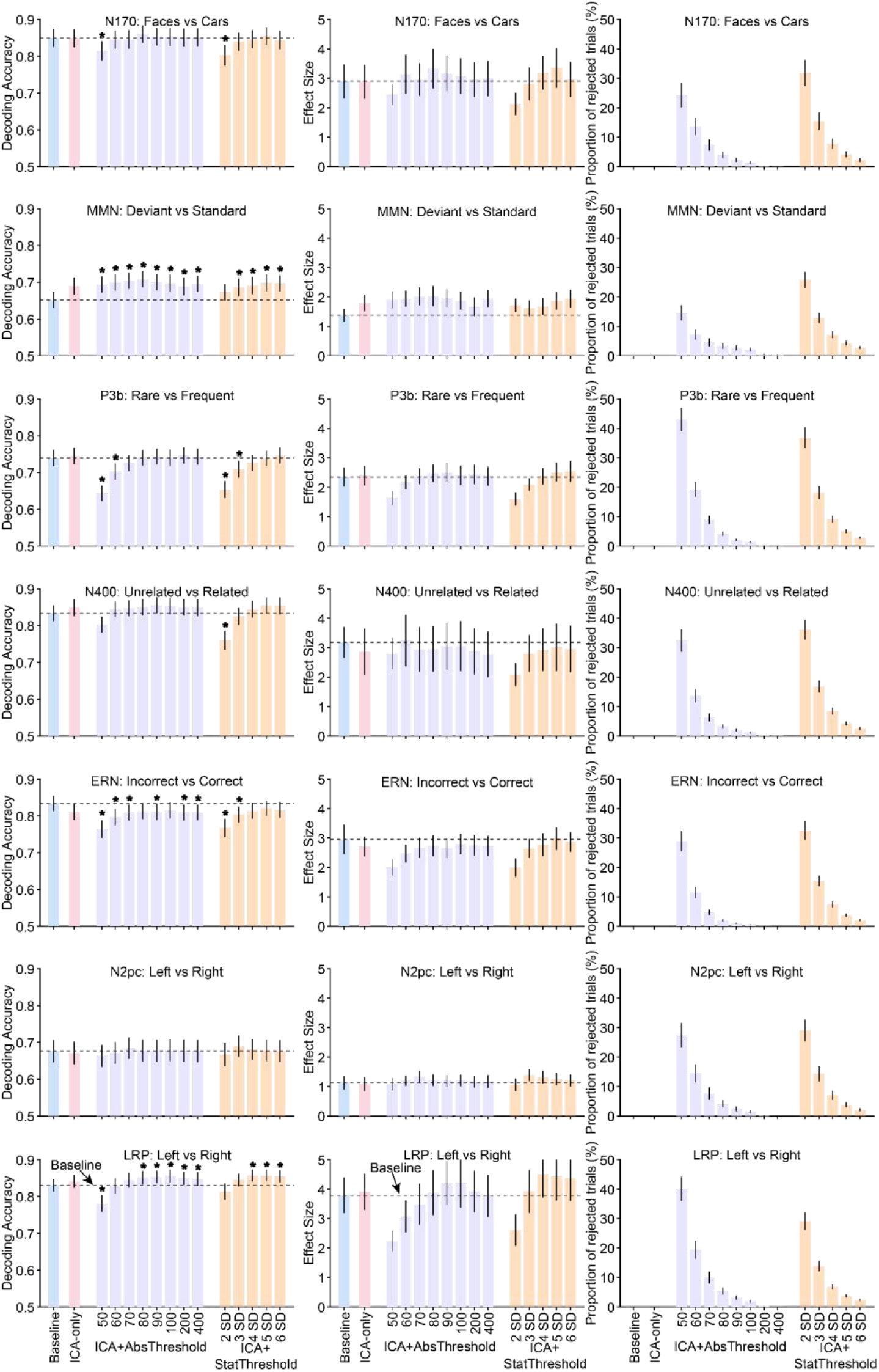
Decoding accuracy (left column), effect size (Cohen’s *d_z_*, middle column), and percentage of rejected trials (right column) resulting from the different artifact minimization approaches for each ERP CORE component, averaged across the time-window shown in Table 1. The different bars within each panel show the results for a given artifact approach and set of parameters. Error bars show the SEM. The dashed horizontal lines show decoding performance in the baseline condition. Asterisks mark cases in which decoding accuracy differed significantly between a given approach and the baseline approach (according to a paired *t* test).

Now we will consider the effects of combining ICA with artifact rejection using several different absolute rejection thresholds (ICA+AbsThreshold). The proportion of trials rejected for each threshold is shown in the right column of Figure 2 (violet bars). As expected, the proportion of rejected trials increased as the threshold decreased, with between 25% and 45% of trials rejected at the lowest threshold. As shown by the violet bars in the left and middle columns of Figure 2, mean decoding accuracy and *d_z_* were slightly but significantly increased by this combination of artifact correction and rejection for the MMN at all thresholds and for the LRP at the higher thresholds. Decoding accuracy and *d_z_* tended to drop when the threshold was reduced all the way down to 50 or 60 µV, yielding significantly worse decoding accuracy relative to baseline for the P3b and LRP. For the ERN, the ICA+AbsThreshold approach yielded a small but statistically significant reduction in decoding accuracy relative to the Baseline approach (but, as described in the Discussion, this reduction may reflect a desirable elimination of blink-related confounds). The same general pattern was observed for the ICA+StatThreshold approach (orange bars).

Thus, for the seven components in the ERP CORE, artifact correction and rejection had only a minimal impact on decoding accuracy unless a very large proportion of trials was rejected. For the threshold levels typically used in ERP research (e.g., 100 µV), the reduction in available trials resulting from artifact rejection was approximately balanced by a reduction in noise, leading to either no change or a small increase in decoding performance (or a small decrease in the case of the ERN).

### 3.2 Challenging multiclass decoding cases

We also investigated more challenging decoding cases in which the stimulus classes being decoded were more similar (i.e., slightly different orientations or locations) and the number of classes was greater (i.e., 12 or 16 classes). In addition, we included a noisier dataset from a much broader sample of participants (Orientations-Community). As shown in Supplementary Table S1, the noise level was almost twice as high for the Orientations-Community participants as for the Orientations-1 and-2 participants.

Figure 3 summarizes the orientation decoding results for the Orientations-1 and Orientations-2 datasets, and Figure 4 summarizes the location decoding results for the Orientations-2 dataset and the orientation decoding results for the Orientations-Community dataset. For each case, the data are summarized for a perceptual processing period (100-500 ms) and a working memory maintenance period (500-1000 ms).

**Figure 3.**
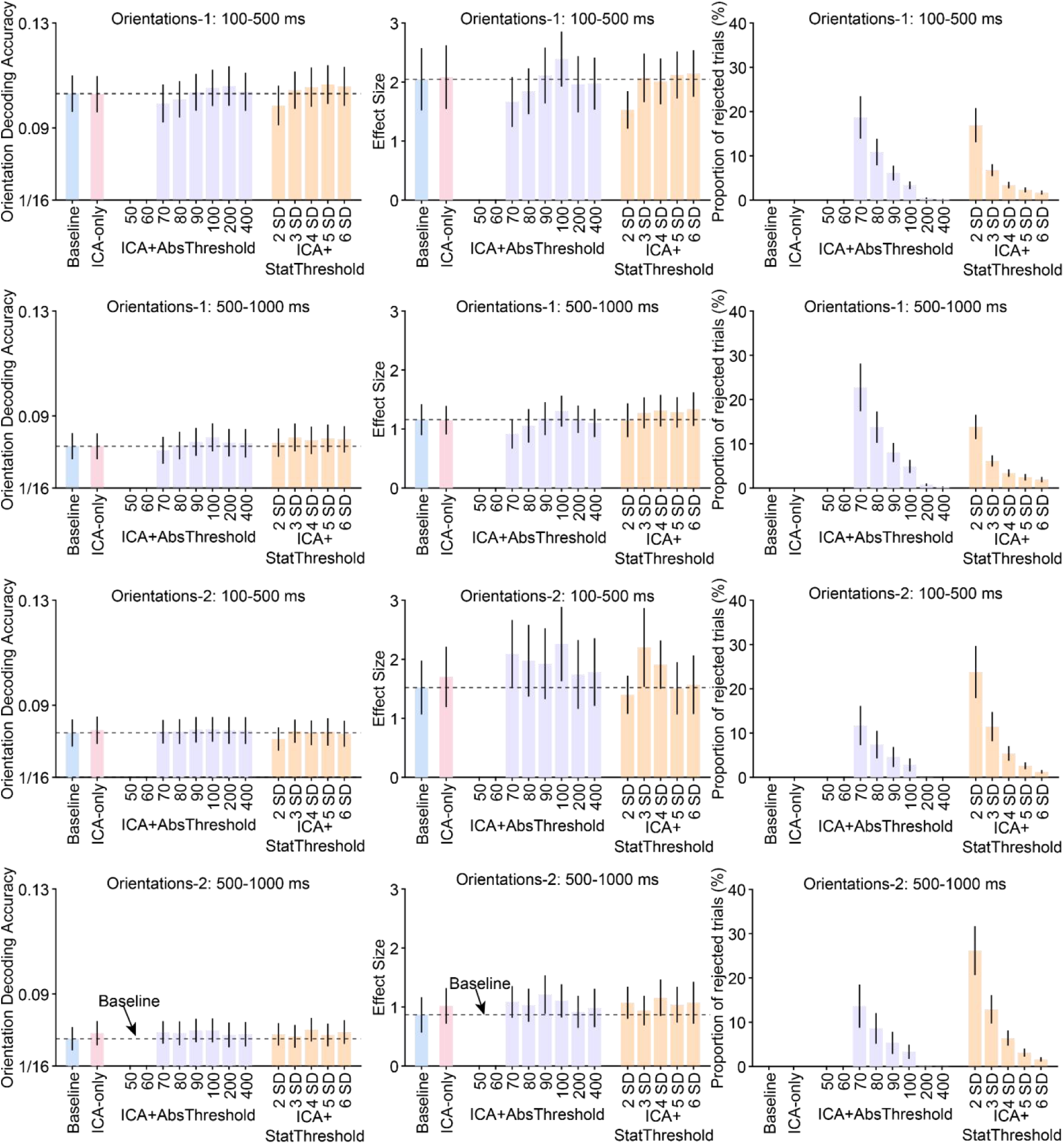
Decoding accuracy (left column), effect size (Cohen’s *d_z_*, middle column), and percentage of rejected trials (right column) resulting from the different artifact minimization approaches for the Orientations-1 and Orientations-2 datasets, separately for a perception time-window (100-500 ms) and a working memory maintenance time window (500-1000 ms). The different bars within each panel show the results for a given artifact approach and set of parameters. Error bars show the SEM. The dashed horizontal lines show decoding performance in the baseline condition. There were no cases in which decoding accuracy differed significantly between a given approach and the baseline approach (according to a paired *t* test).

**Figure 4.**
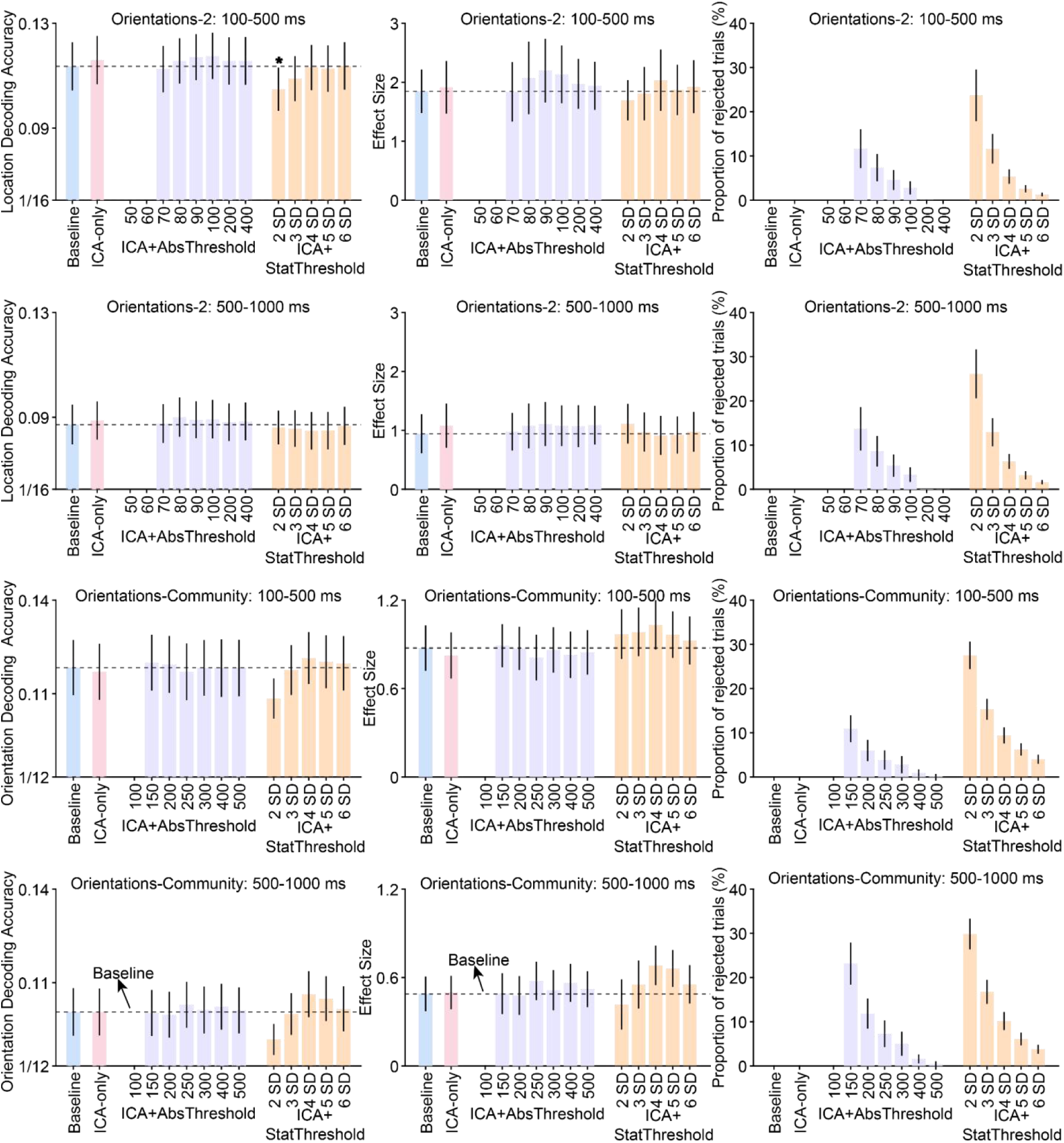
Decoding accuracy (left column), effect size (Cohen’s *d_z_*, middle column), and percentage of rejected trials (right column) resulting from the different artifact minimization approaches for decoding of stimulus location in the Orientations-2 dataset and decoding of orientation in the Orientations-Community dataset, separately for a perception time-window (100-500 ms) and a working memory maintenance time window (500-1000 ms). The different bars within each panel show the results for a given artifact approach and set of parameters. Error bars show the SEM. The dashed horizontal lines show decoding performance in the baseline condition. An asterisk marks the one case in which decoding accuracy differed significantly between a given approach and the baseline approach (according to a paired t test).

In all of these cases, ICA had little or no impact: decoding accuracy and *d_z_* were approximately the same for the ICA-Only approach as for the Baseline approach, and only one of the differences was statistically significant (a significant reduction in decoding accuracy for the strictest ICA+StatThreshold threshold in the upper left panel of Figure 4). Adding artifact rejection to ICA also had little or no impact: decoding accuracy and *d_z_* were approximately the same for the ICA+AbsThreshold and ICA+StatThreshold approaches as for the ICA-Only and Baseline approaches. Decoding performance dropped off slightly for the lowest thresholds in the Orientations-1 dataset, but these differences were modest and not statistically significant.

## 4. Discussion

In this study, we investigated whether several artifact minimization approaches enhanced the performance of SVM-based decoding analyses across a wide range of EEG/ERP paradigms. We have previously assessed the effects of these approaches on conventional univariate ERP analyses (Zhang, Garrett, Simmons, et al., 2024). However, this is the first time that a broad range of datasets has been used to assess the impact of artifact correction and artifact rejection on decoding performance. We found that none of the artifact minimization approaches consistently improved decoding accuracy or effect sizes across a broad range of datasets.

ICA-based correction might have been expected to increase decoding accuracy by reducing large voltage deflections arising from blinks and eye movements, but it did not significantly improve decoding accuracy in any of the 15 datasets we examined. It is possible, however, that correction of blinks and eye movements would improve decoding for effects that are largest near the eyes, where the uncorrected voltage deflections would be the largest.

The rejection of trials with large voltage deflections led to a significant reduction in decoding accuracy for several of the ERP CORE datasets when very strict rejection thresholds were used. In these cases, the cost of the smaller number of trials for training the decoder presumably outweighed any benefits from noise reduction. These strict thresholds were also problematic for the other datasets, where so many trials were rejected that decoding could not even be performed for some of the participants. For more typical rejection thresholds, the combination of ICA and artifact rejection led to a small but significant increase in decoding accuracy in two of the 15 datasets (MMN and LRP) and no significant change in the other datasets. It is not clear whether there is something special about the two datasets with significant effects that made artifact minimization more beneficial or whether these were spurious or idiosyncratic effects.

The combination of ICA and artifact rejection actually led to a small but significant decrease in decoding accuracy in the ERN dataset across a broad range of rejection thresholds. For the ERN, we have previously shown that blinks differ systematically between error trials and correct trials, inflating the difference in voltage between these trial types (Zhang, Garrett, Simmons, et al., 2024). These blinks may have artificially inflated decoding accuracy in the Baseline condition, and the artifact correction and rejection procedures may have reduced this inflation. This was also observed in other research, where the decoding accuracy for anterior frontal channels was significantly but artifactually larger in uncorrected data compared to corrected data (Hong et al., 2020).

Thus, artifact correction and rejection may be valuable for reducing confounds from non-neural artifacts even if they produce a slight reduction in decoding accuracy.

The general conclusion that can be drawn from the entire set of results is that the artifact minimization approaches tested here have little or no impact on decoding accuracy (assuming that rejection thresholds are not too strict). This is consistent with prior studies that examined a more restricted range of datasets (Bibián et al., 2022; Kang et al., 2024).

This is generally good news, because it is one less thing for researchers to worry about, assuming that their datasets are not too different from the 15 datasets examined here.

However, we would like to make two recommendations. First, we recommend that researchers perform artifact correction to minimize the voltages associated with blinks and eye movements. This is because these artifacts may differ systematically across conditions (Hong et al., 2020; Zhang, Garrett, Simmons, et al., 2024), and these non-neural signals may artificially inflate decoding accuracy if not corrected. There is no obvious downside to correction of ocular artifacts in decoding analyses, and the elimination of confounds is a clear upside. Our second recommendation is that if researchers choose to reject trials with large voltage deflections, they should not use overly strict thresholds (e.g., less than 100 µV or less than 3 SD). Rather than being helpful, strict thresholds may reduce decoding accuracy and effect sizes.

These recommendations should be valid across a fairly broad range of situations given that we observed fairly similar results across a broad range of datasets. These datasets varied in the nature of the information being decoded, the number of participants, the number of experimental stimuli, the number of electrode sites, and the density of the electrode arrays. However, these datasets were all from neurotypical adults, and the present findings may not generalize to data recorded from other types of populations, such as infants and people with psychiatric and neurological disorders (Ashton et al., 2022; Ng et al., 2022; Rashid & Calhoun, 2020). In addition, all of the datasets examined here were recorded under laboratory conditions using very high quality recording systems, and the results may not generalize to datasets collected with different recording systems such as dry electrodes (G.-L. Li et al., 2020; Shad et al., 2020). Different results may also be found for mobile EEG experiments, which may require higher rejection thresholds to achieve optimal decoding performance (Biondi et al., 2022; Lau-Zhu et al., 2019; Richer et al., 2024).

It is also possible that other artifact minimization approaches would have greater benefits than the approaches tested here. For example, we used very simple approaches to detect extreme voltages, and we applied the same threshold across all participants and electrode sites. Other algorithms for rejecting segments with extreme values may be better (Chang et al., 2020; Jas et al., 2017), as may adaptive thresholding approaches (Talsma, 2008).

Furthermore, our artifact correction procedure used EEGLAB’s default ICA algorithm and parameters, as is common in EEG research. Other advanced approaches, such as deep learning-based methods (Yang et al., 2018), probability mapping algorithms (Islam et al., 2021), and variants of ICA (Mammone et al., 2012; Sai et al., 2018; Stergiadis et al., 2022; Sun et al., 2021), may correct artifacts more efficiently. Similarly, the present findings may not generalize to more advanced decoding algorithms, such as convolutional neural networks, recurrent neural networks, and deep belief networks for deep learning (Al-Saegh et al., 2021; Craik et al., 2019; Gong et al., 2022). There is a large universe of possible artifact minimization and decoding approaches, and a wide-ranging examination is beyond the scope of the present study. However, now that we have identified a broad range of datasets and developed a straightforward approach for assessing the effectiveness of artifact minimization approaches, future research can determine whether artifact minimization approaches exist that produce a large improvement in decoding accuracy.

## CRediT authorship contribution statement

**Guanghui Zhang**: Writing – review & editing, Writing – original draft, Visualization, Validation, Supervision, Software, Resources, Methodology, Investigation, Conceptualization. **Steven J. Luck**: Writing – review & editing, Writing – original draft, Visualization, Validation, Supervision, Software, Project administration, Methodology, Funding acquisition, Conceptualization.

## Acknowledgments

For their thoughtful insights, we thank the members of the Luck Lab, including but not limited to: Carlos Carrasco, John Kiat, David Garrett, Kurt Winsler, and Brett Bahle. We also thank Gi-Yeul Bae from Arizona State University for sharing his datasets with us.

---

**Figure S1.**
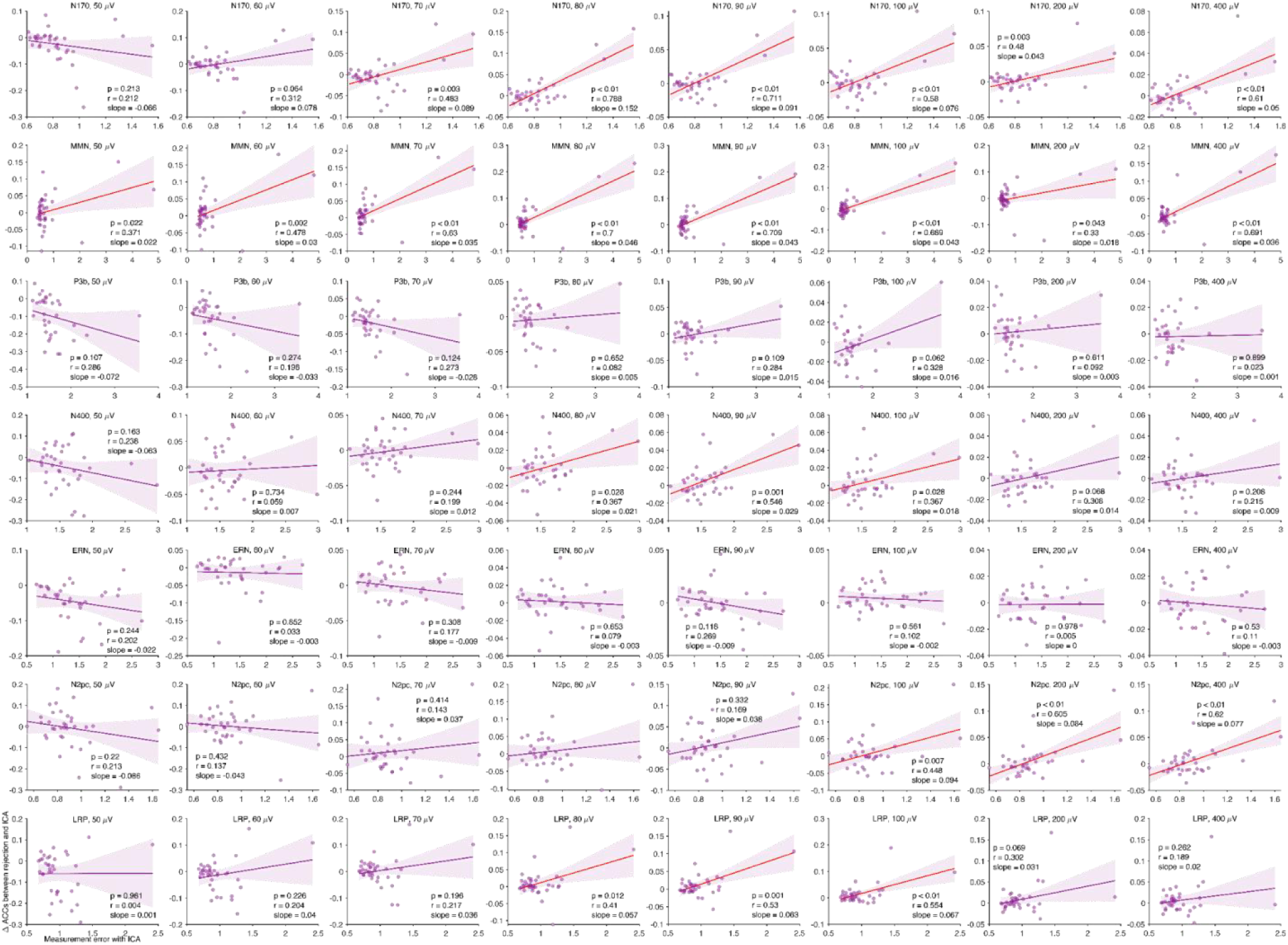
Scatterplots of the relationship between measurement error (X axis) and the difference in decoding accuracy between the ICA+AbsThreshold approach and the ICA-Only approach (Y axis) for different rejection thresholds and each of the ERP components from the ERP CORE. Note that we used the simple voltage threshold (SVT) algorithm and moving window peak-to-peak (MWP) algorithm in ERPLAB to detect and reject trials with artifacts with thresholds of 50, 60, 70, 80, 90, 100, 200, or 400 µV. To estimate the measurement error, we first computed the standard error of the mean (SEM) of the amplitudes across trials at each time point within the measured time window (see Table 1 for time window for each component) for each experimental condition in a given participant, and we then calculated the root mean square of these SEM values.

**Figure S2.**
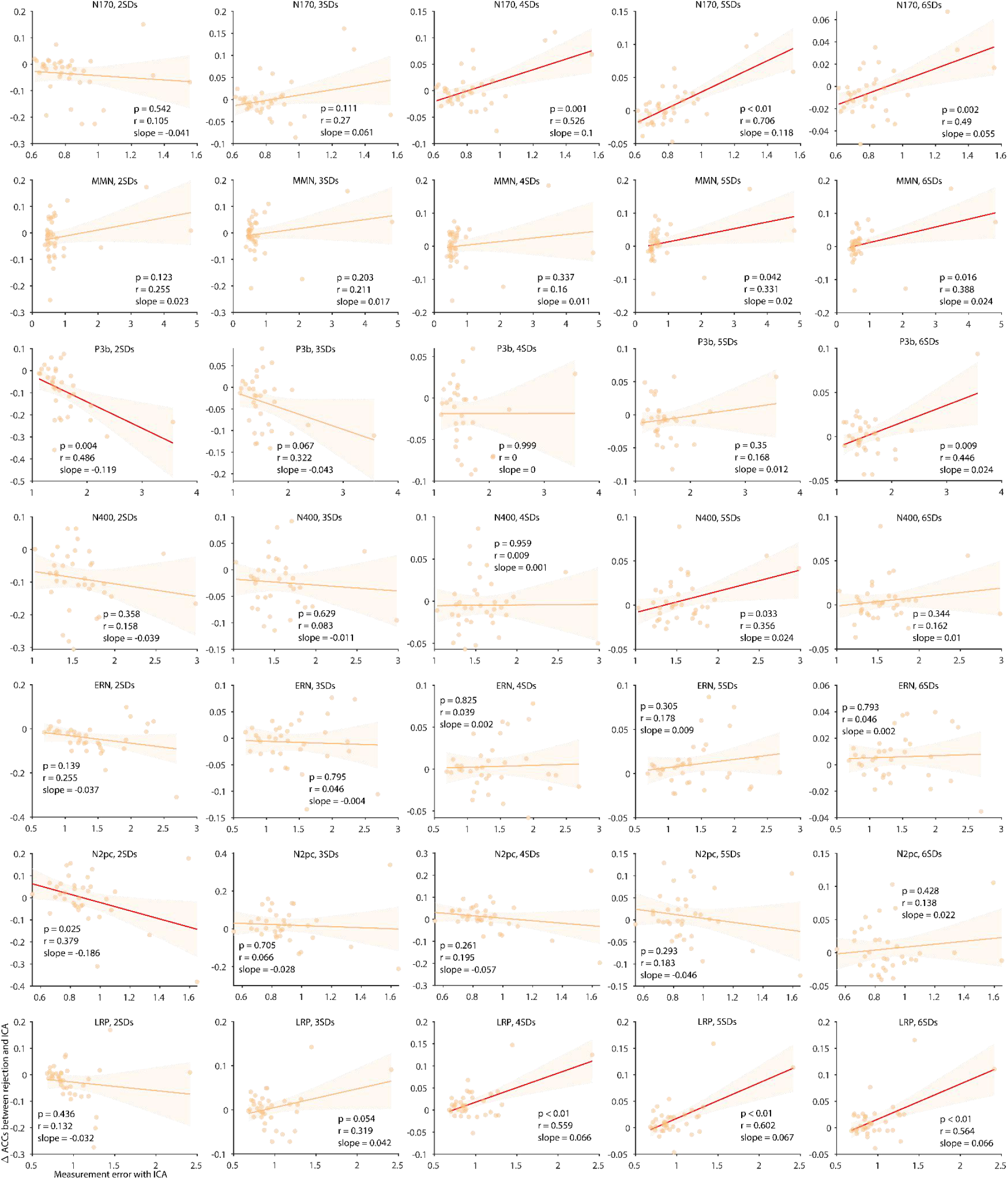
Scatterplots of the relationship between measurement error (X axis) and the difference in decoding accuracy between the ICA+StatThreshold approach and the ICA-Only approach (Y axis) for different rejection thresholds and each of the ERP components from ERP CORE. Note that we used the probability density algorithm to detect and reject trials with artifacts with thresholds of 2,3, 4, 5, or 6 standard deviations (SDs) from the mean.

**Figure S3.**
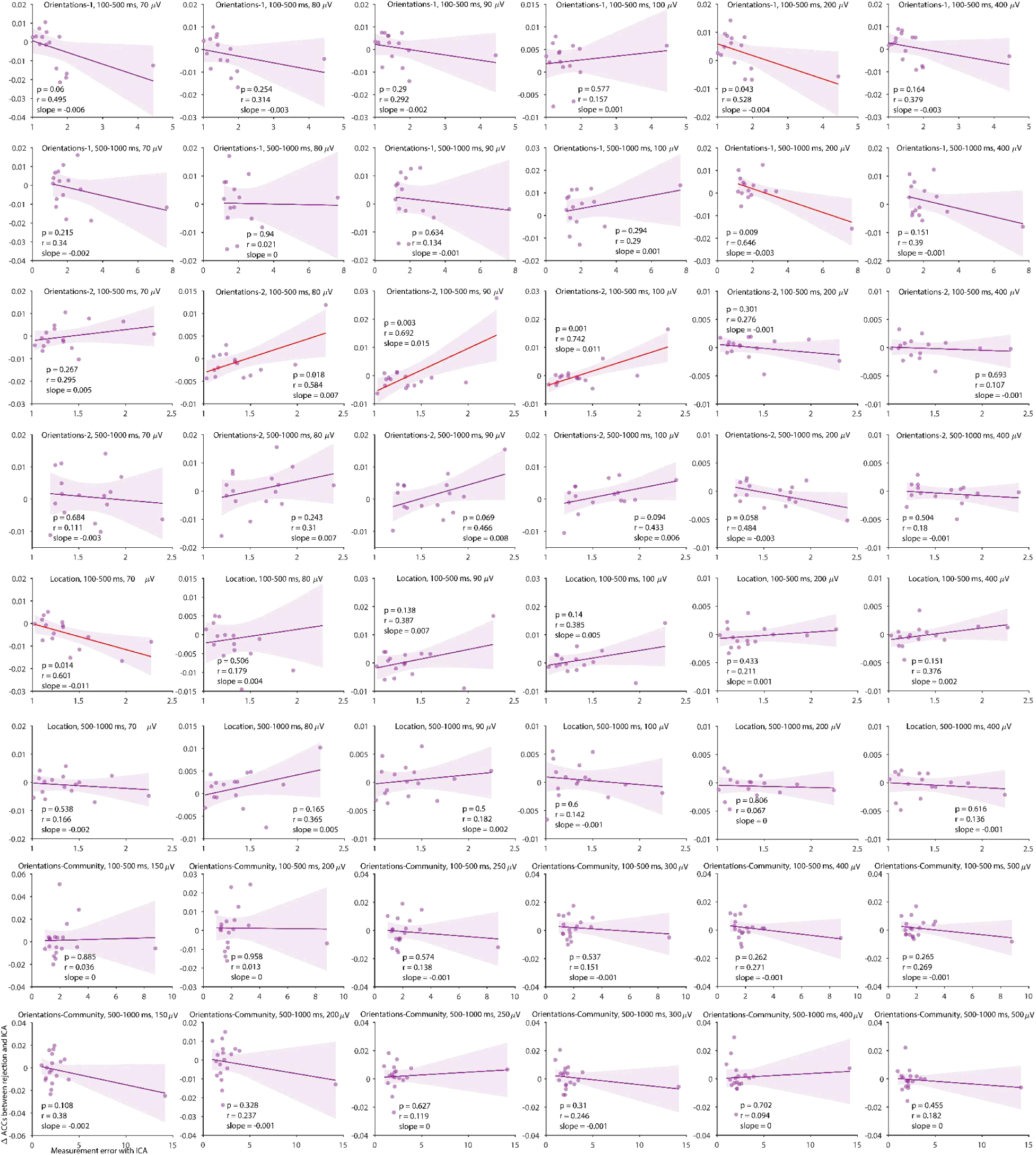
Scatterplots of the relationship between measurement error (X axis) and the difference in decoding accuracy between the ICA+AbsThreshold approach and the ICA-Only approach (Y axis) for different rejection thresholds and each of the ERP components from the Orientations-1, Orientations-2, and Orientations-Community experiments. Note that we used the simple voltage threshold (SVT) algorithm and moving window peak-to-peak (MWP) algorithm in ERPLAB to detect and reject trials with artifacts with thresholds of 50, 60, 70, 80, 90, 100, 200, or 400 µV.

**Figure S4.**
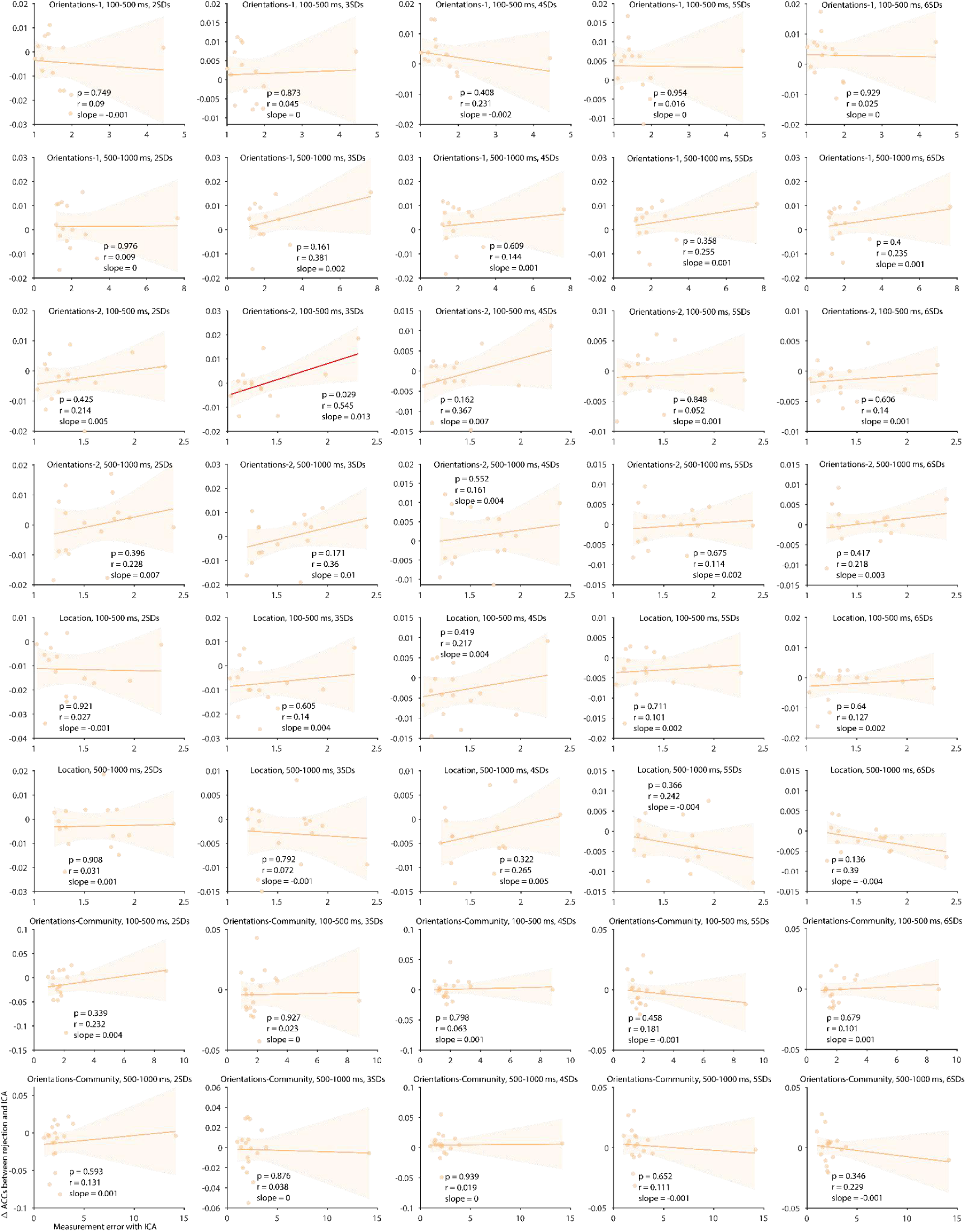
Scatterplots of the relationship between measurement error (X axis) and the difference in decoding accuracy between the ICA+StatThreshold approach and the ICA-Only approach (Y axis) for different rejection thresholds and each of the ERP components from the Orientations-1, Orientations-2, and Orientations-Community experiments. Note that we used the probability density algorithm to detect and reject trials with artifacts with thresholds of 2, 3, 4, 5, or 6 standard deviations (SDs) from the mean.

**Table S1.**
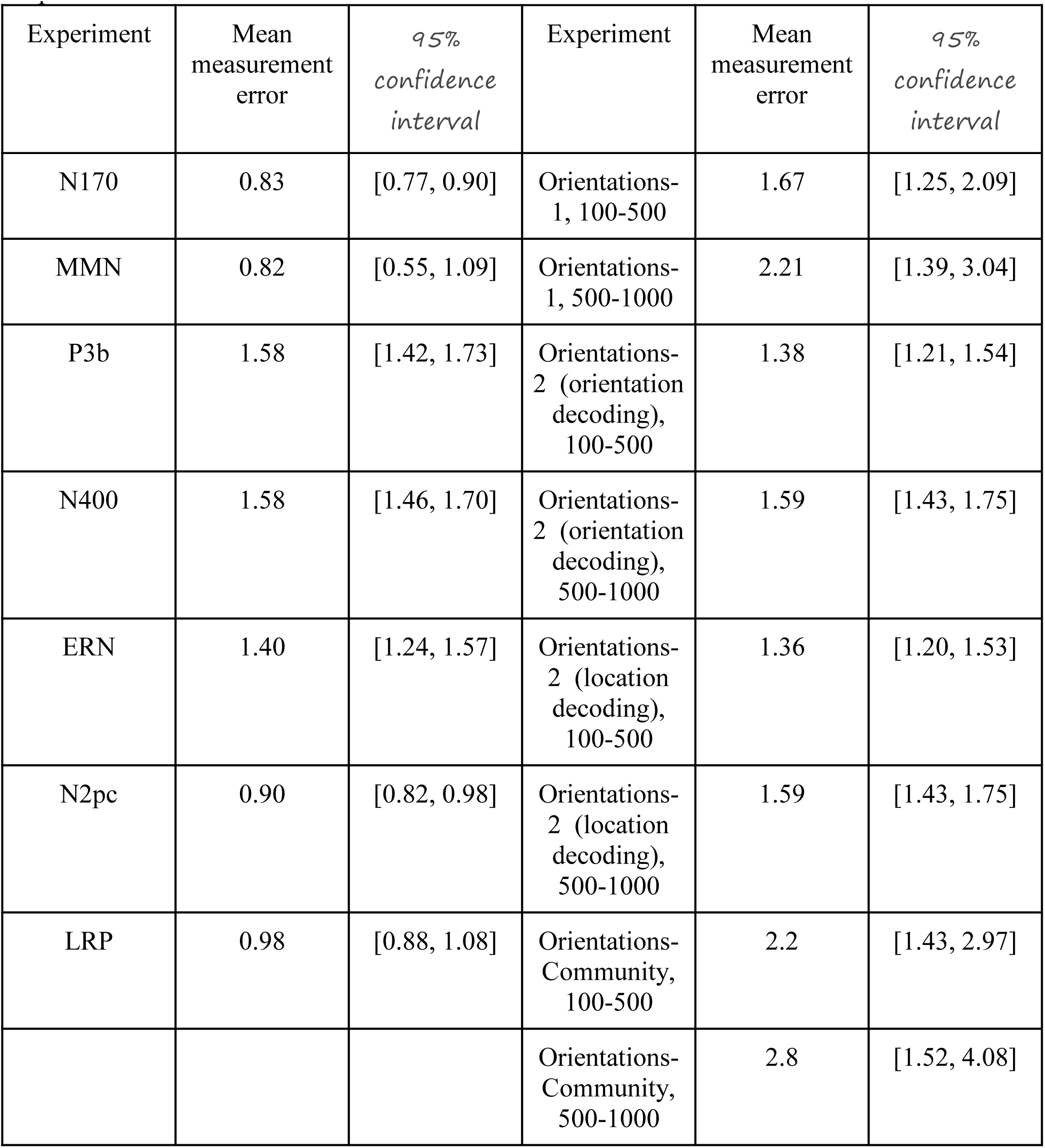
Mean and 95% confidence interval of the measurement error values for different experiments.

## Footnotes

1 Although this narrow bandpass can significantly distort the time course of the ERP waveform (Zhang, Garrett, & Luck, 2024a, 2024b), it does not alter scalp distributions, thus preserving the integrity of the ICA decomposition process.

